# Elucidation of neuronal activity in mouse models of TMJ injury by *in vivo* GCaMP Ca*^2+^* imaging of intact trigeminal ganglion neurons

**DOI:** 10.1101/2024.01.16.575919

**Authors:** Hyeonwi Son, John Shannonhouse, Yan Zhang, Ruben Gomez, Man-Kyo Chung, Yu Shin Kim

**Author notes:** Corresponding author: Yu Shin Kim, PhD, Department of Oral & Maxillofacial Surgery, University of Texas Health Science Center at San Antonio 7703 Floyd Curl Drive, San Antonio, TX 78229.

## Abstract

Patients with temporomandibular disorders (TMD) typically experience facial pain and discomfort or tenderness in the temporomandibular joint (TMJ), causing disability in daily life. Unfortunately, existing treatments for TMD are not always effective, creating a need for more advanced, mechanism-based therapies. In this study, we used *in vivo* GCaMP3 Ca^2+^ imaging of intact trigeminal ganglia (TG) to characterize functional activity of the TG neurons *in vivo*, specifically in TMJ animal models. This system allows us to observe neuronal activity in intact anatomical, physiological, and clinical conditions and to assess neuronal function and response to various stimuli. We observed a significant increase in spontaneously and transiently activated neurons responding to mechanical, thermal, and chemical stimuli in the TG of forced mouth open (FMO) mice. An inhibitor of the CGRP receptor significantly attenuated FMO-induced facial hypersensitivity. In addition, we confirmed the attenuating effect of CGRP antagonist on FMO-induced sensitization by *in vivo* GCaMP3 Ca^2+^ imaging of intact TG. Our results contribute to unraveling the role and activity of TG neurons in the TMJ pain animal models of TMD, bringing us closer understanding the pathophysiological processes underlying TMD. Our study also illustrates the utility of *in vivo* GCaMP3 Ca^2+^ imaging of intact TG for studies aimed at developing more targeted and effective treatments for TMD.

## Introduction

Joints endow us with flexible movements like moving away from danger, securing food, intricate communication, and manipulating food or objects. Among those joints, the temporomandibular joint (TMJ) performs essential behaviors such as mastication, speech, and other jaw movement activities. The paramount role of joints is evident as 80% of joint afferent nerve fibers exhibit nociceptive properties[23]. The dominance of nociceptors underscores their critical function—specialized nerve endings highly sensitive to potential damage & injury stimuli. Their prevalence indicates an evolutionary response toward immediate detection and mitigation of injury, serving a protective role for joints[34].

The protective mechanism provided by joint afferents can become a double-edged sword when injured and inflamed. For instance, excessive mechanical stress can induce disc displacement in TMJ, leading to inflammation[30]. Nociceptors could also cause a prolonged increase in excitability after inflammation and become more sensitive to normally innocuous stimuli[6]. This peripheral sensitization in TMJ afferents, innervated exclusively by trigeminal ganglion neurons, results in temporomandibular disorder (TMD)-related pain and discomfort[6,31]. As a result, the pain and discomfort occurring as a protective response to prevent further injury can become persistent, even in the absence of an immediate threat to the joint’s integrity[8].

TMD is the second most common musculoskeletal disorder, affecting 5-12% of the US population (NIH). It consists of myalgia, arthralgia, and myofascial pain, resulting in headaches, limited jaw motion, and pain-related disability[9]. Existing treatments for TMD are not always effective in all patients, underscoring the need for more advanced, mechanism-based therapies. Therapeutic limitations stem from our poor understanding of TMD mechanisms. Since TMD pain exclusively relies on TG neurons relaying pain signals from the periphery to the central nervous system[8], monitoring the activities of TG neurons could provide direct mechanistic information enabling identification of the pathological mechanisms of TMD. We have here confirmed peripheral sensitization in TMD animal models using *in vivo* GCaMP3 Ca^2+^imaging of intact TG and have demonstrated the utility of GCaMP3 Ca^2+^ imaging for unraveling the function and activity pattern of TG neurons in the TMD animal model. Our study shows that *in vivo* GCaMP3 Ca^2+^ imaging of intact TG offers unprecedented insights into the functional alterations of TG neurons in TMD animal models. Thus, our approach promises to bridge existing knowledge gaps and foster the development of targeted therapeutic interventions.

## Methods

### Animals

*Pirt*-GCaMP3 mice, prepared according to methods described previously[18,19], were used with C57BL/6 mice. Mice were housed 3-5 mice per cage and exposed to a 12-hour light/dark cycle. They were allowed access to water and mouse chow *ad libitum*. Male and female mice, aged between 6-12 weeks, were used for experiments. All animal procedures were approved by the University of Texas Health at San Antonio (UTHSA) Animal Care and Use Committee (IACUC). All experiments were performed following the National Institute of Health Guide for the Care and Use of Laboratory Animals.

### Peptides and drugs

α-CGRP was acquired from Bachem (Bubendorf, Switzerland). CFA was sourced from Sigma (Burlingame, CA, USA). BIBN-4096 was obtained from Tocris Bioscience/Bio-Techne Corporation (Minneapolis, MN, USA). BIBN-4096, at a concentration of 10 mg/kg, was formulated in a solution containing 5% DMSO and 5% Tween-80 dissolved in 0.9% NaCl. The prepared solution was administered intraperitoneally to the experimental subjects.

### CFA Injection

Mice of the C57BL/6 strain were subjected to unilateral injections into the Temporomandibular Joint (TMJ) intra-articular space using Complete Freund’s Adjuvant (CFA). CFA was emulsified 1:1 with PBS. Depending on the experimental design, these injections are administered once or twice. As a control measure, saline was injected on the contralateral side.

### FMO (Forced Mouth Opening)

Mice were anesthetized with Ketamine/Xylazine (80/10 mg/kg). While anesthetized, the mouth was opened using a colibri retractor (Fine science tools, 17000-03), and remained at its maximum opening for 3 hours each day over five consecutive days. Control mice underwent the same anesthetic regimen without a sustained mouth-opening procedure.

### Facial von Frey test

To ensure consistent behavioral responses, mice underwent a habituation phase in which they were familiarized with the experimenter’s scent and hand touch for 2 days. The mice were then acclimated to a transparent plexiglass chamber equipped with 4 oz paper cups for 2 hours daily over 3 consecutive days. After acclimation, baseline tests were carried out to assess the cutaneous sensitivity of the facial area, particularly the temporomandibular region, using von Frey filaments, for 5-7 days. The facial von Frey test baseline was determined to be achieved when mice exhibited a threshold between 0.5 and 0.7 g. The exact thresholds were ascertained employing the Dixon “up-and-down” methodology[14].

### Trigeminal ganglion (TG) exposure surgery for *in vivo Pirt*-GCaMP3 Ca^2+^ imaging

Mice were anesthetized by i.p. injection of Ketamine (Zoetis, Parsippany-Troy Hills, NJ, USA)/Xylazine (approximately 80/10 mg/kg) (VetOne, Boise, ID, USA), and ophthalmic ointment (Lacri-lube; Allergen Pharmaceuticals) was applied to the eyes. The right-side dorsolateral skull was exposed by removing skin and muscle. A patch of dorsolateral skull (parietal bone between the right eye and ear) was removed using a dental drill (Buffalo Dental Manufacturing, Syosset, NY, USA) to make a cranial window (∼10X10 mm). The TG was then exposed by aspirating overlying cortical tissue. During the surgery, the mouse’s body temperature was maintained on a heating pad at 37°C ± 0.5°C and monitored by a rectal probe.

### *In vivo Pirt*-GCaMP3 Ca^2+^ imaging of intact TG

*In vivo* Pirt-GCaMP3 Ca^2+^ imaging of intact TG in live mice was performed for 2-5 hr immediately after exposure surgery. After the exposure surgery, mice were placed abdomen-down on a custom-designed platform under the microscope. For *in vivo* Pirt-GCaMP3 Ca^2+^ imaging of intact TG, the animal’s head was fixed by a head holder to minimize movements from breathing and heartbeats. During the imaging session, body temperature was maintained at 37°C ± 0.5°C on a heating pad and was monitored by a rectal probe. Anesthesia was maintained with 1-2% isoflurane using a gas vaporizer with pure oxygen. Live images were acquired at ten frames per cycle in frame-scan mode at ∼4.5 to 8.79 s/frame, ranging from 0 to 900 µm, using a 5 × 0.25 NA dry objective at 512 × 512 pixels or higher resolution with solid diode lasers tuned at 488 nm and emission at 500-550 nm. von Frey filaments (0.4 g) and noxious water (4°C and 50°C) were applied to the different divisions of the animal’s face divided by TG branches. Capsaicin (500 mM, 10 μl) was injected intracutaneously into the different TG branches.

For the analysis of imaging data, raw image stacks were collected, deconvoluted, and imported into ImageJ (NIH). We realigned and corrected optical planes from sequential time points utilizing the stackreg plugin, which is based on rigid-body cross-correlation image alignment. We expressed Ca^2+^ signal amplitudes as a ratio of F_t_ (the fluorescence intensity in each frame) to F_0_ (the average fluorescence intensity observed during the initial one to four frames). Each responding cell was confirmed through a meticulous visual examination of the raw imaging data.

### Statistics

Statistical evaluations were conducted using Prism (GraphPad). Error bars represent mean ± S.E.M. A p<0.05 was considered significant. The specific statistical tests employed are detailed in the figure legends.

## Results

### *In vivo* GCaMP Ca^2+^ imaging enables monitoring of functional changes and activity patterns of TG neurons innervating temporomandibular region (V3)

Using *Pirt*-GCaMP3 mice, in which the genetically encoded Ca^2+^ indicator GCaMP3 is specifically expressed in >95% of all peripheral sensory neurons under the control of the *Pirt* Promoter[18,19], we imaged intact TG *in vivo* (Fig. 1A). We imaged 2819±18.84 neuronal cell bodies in each TG examined. Diameters of most neurons ranged from 10 to 35 μm (Fig. 1B), which we classified into three groups, including small (<20 μm), medium (20-25 μm), and large (>25 μm) diameter neurons. Small diameter TG neurons were the most numerous (Figs. 1C and D). To observe the basal condition of spontaneously activated neurons, prior to stimulus or drug application, we monitored TG neurons for approximately 25 minutes via *in vivo* Ca^2+^ imaging of intact TG. An average of 36.3 neurons exhibited spontaneous activity (Figs. 1E and F). Small-diameter TG neurons exhibited the highest average activation, followed by medium diameter neurons (Figs. 1G and H).

**Figure 1.**
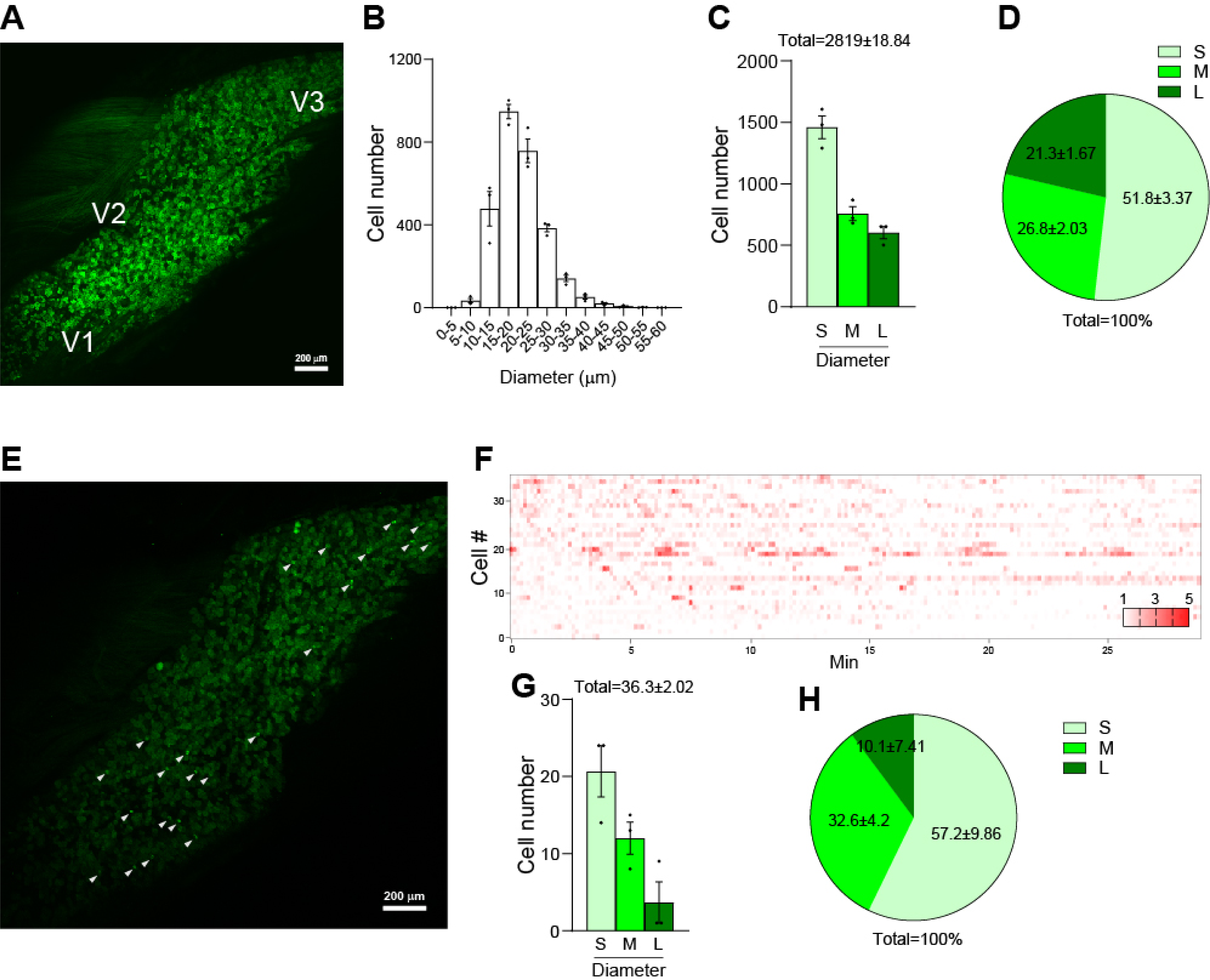
Visualization and Analysis of Activated Neuronal Populations in Trigeminal Ganglion of *Pirt*-GCaMP3 Mice. **(A)** Representative image of whole TG captured via *in vivo Pirt*-GCaMP3 Ca^2+^ imaging of intact TG. The locations of neuronal cell bodies for V1 (ophthalmic), V2 (maxillary), and V3 (mandibular) are indicated. (**B**) Histogram depicting the cell number distribution according to cell diameter (μm) (n = 3 mice). (**C**) Graph showing the cell numbers categorized into three groups based on cell diameter: small (<20 μm), medium (20-25 μm), and large (>25 μm) (n = 3 mice). (**D**) Pie chart representing the percentage composition of each cell diameter size group (n = 3 mice). (**E**) Representative image of spontaneous activities captured through *in vivo Pirt*-GCaMP3 Ca ^2+^ imaging of intact TG. White arrowheads point to spontaneously activated neurons. (**F**) Representative heatmap displaying the spontaneously activated individual neurons in normal mice. (**G**) Bar graph showing the number of spontaneously activated neurons, categorized by cell diameter size group (n = 3 mice). (**H**) Pie chart showing the percentage composition of spontaneously activated neurons according to cell diameter size (n = 3 mice). S: small diameter TG neurons (<20 μm); M: medium (20-25 μm); L: large (>25 μm). Error bars indicate S.E.M.

Next, to observe TG neuronal activity in response to various stimuli, we applied von Frey filament (mechanical), hot/cold water (thermal), or capsaicin (chemical) to each orofacial region innervated by ophthalmic (V1), maxillary (V2), and temporomandibular (V3) TG neurons[10] during *in vivo Pirt*-GCaMP3 Ca^2+^ imaging of intact TG (Figure 2A). In response to stimuli applied to V1, V2, and V3 regions of the face, TG neurons exhibited a region-specific activation pattern, highlighting a distinct, localized response in the TG neurons associated with each peripheral tissue (Fig. 2A, right panels). V2 and V3 neurons were more responsive to stimuli than V1 neurons (Figs. 2B-E). Neurons activated in response to stimulation in region V3 were primarily medium-diameter (Figs. 2B-E). These results, utilizing *Pirt*-GCaMP3 mice, indicated that *in vivo* GCaMP Ca^2+^ imaging of intact TG in living mice provides a robust tool for capturing neural dynamic responses with various stimuli under various conditions, especially within the V3 region. These techniques and tools will likely provide the capacity for precise and detailed examination of neural signaling and circuit responses.

**Figure 2.**
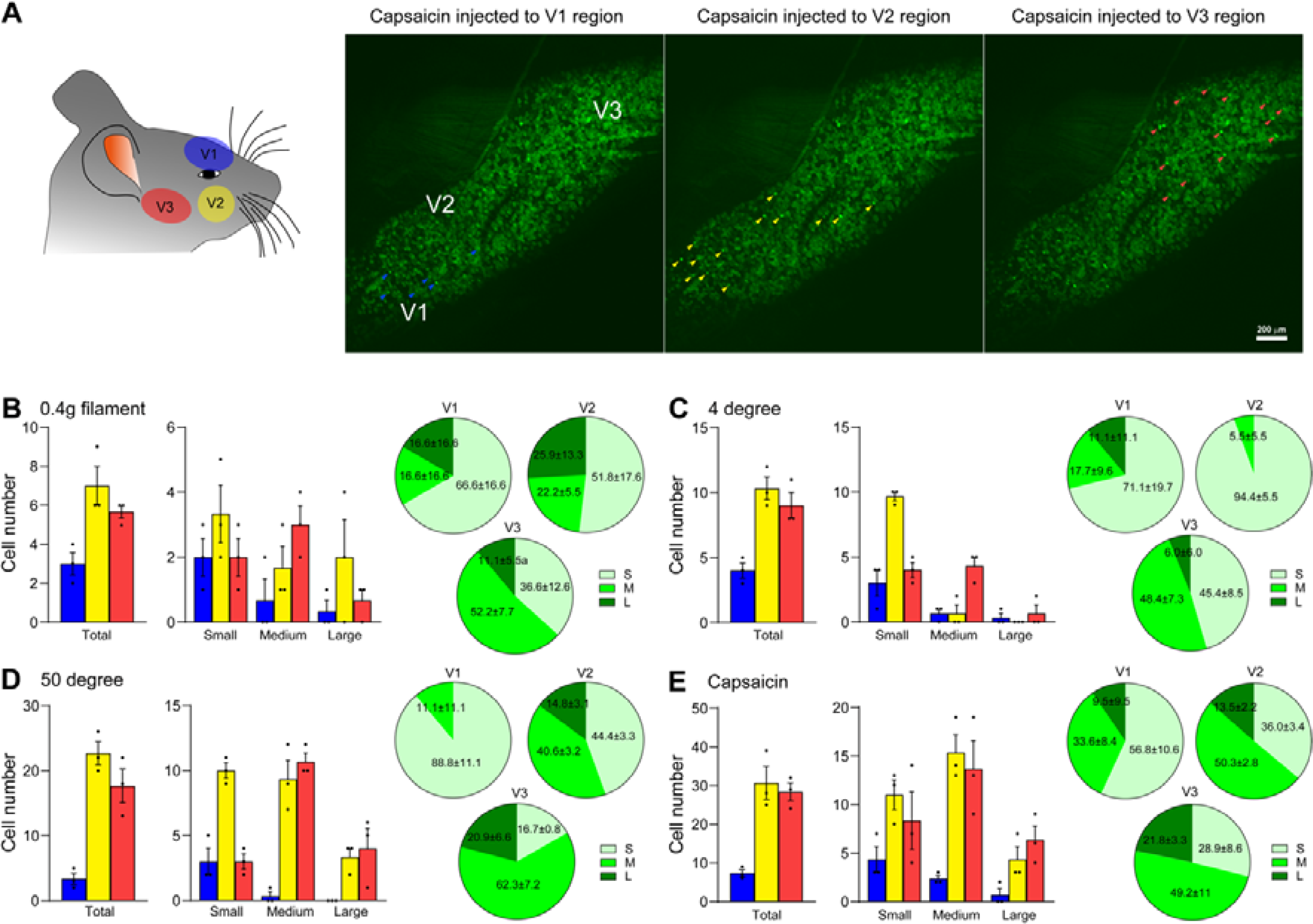
Region-Specific Activation Patterns of TG Neurons in Response to Various Stimuli. **(A)** (*left*) Schematic illustrating the specific regions stimulated during *in vivo* GCaMP3 Ca^2+^ imaging. (*right*) Representative images depicting neurons activated following capsaicin injection in the V1, V2, or V3 regions. (**B-E**) (*left*) Graphs showing the number of individual TG neurons activated by various stimuli: (B) 0.4 g von Frey filament, (C) 4lJ water, (D) 50lJ water, and (E) capsaicin (500 μM, 10 μl) in normal mice (n = 3 mice). Each graph categorizes activated neurons by cell diameter size: small (<20 µm), medium (20-25 µm), and large (>25 µm). (*right*) Pie charts representing the percentage composition of each cell diameter size group (n = 3 mice). Error bars indicate S.E.M.

### Sensitization of TG neurons in TMD animal models

Animal models mimicking the high levels of inflammation observed in the TMJ of TMD patients have been developed using direct injection of chemical agents, including complete Freund’s adjuvant (CFA), zymosan, carrageenan, mustard oil, monoiodoacetate, albumin, and formalin, into the TMJ to cause inflammation. It has been confirmed that intra-TMJ injections of CFA cause mechanical hyperalgesia within the TMJ in rodents[9]. Consistent with this, we confirmed hypersensitivity lasting up to 7 days in the V3 region of CFA-injected mice by the von Frey test (Fig. 3A). Although the current model is essential for deciphering the inflammatory processes leading to TMJ pain and degeneration, there is a pressing need for a direct translational model that accurately replicates the pathological progression of TMD. This necessity stems from the fact that the use of CFA injections only captures some aspects of the complex disease pathogenesis of TMD. The forced mouth opening (FMO) model involves overloading the TMJ and mechanically mimics the pathogenesis of TMD[15,33]. Here, we found that FMO for 3 hours per day for 5 days caused mechanical hypersensitivity in the V3 region, which lasted longer than the CFA-injection model (Fig. 3B). FMO-induced hyperalgesia lasts up to 32 days[33]. These results suggest that, as a translational model for understanding the pathophysiology of TMD, the FMO model is more representative of human TMD.

**Figure 3.**
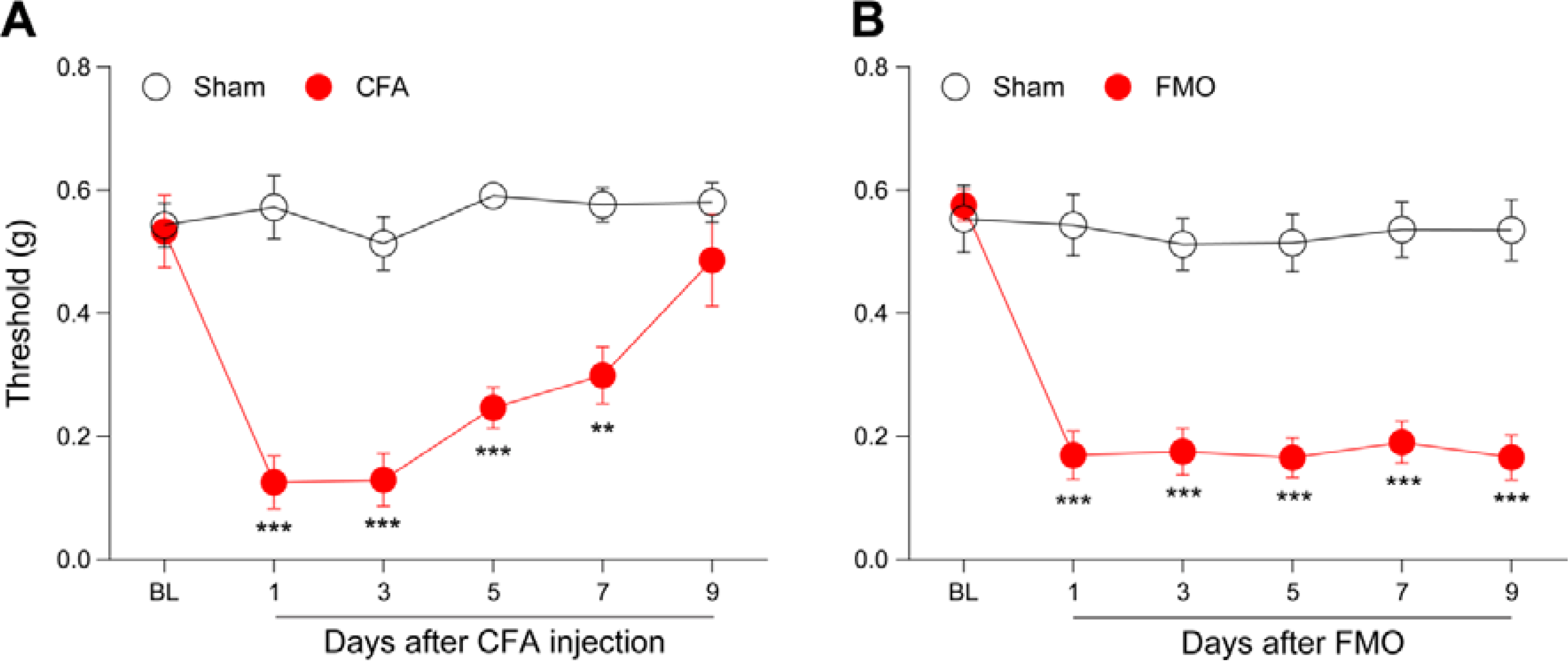
Mechanical Hypersensitivity in the orofacial V3 Region of CFA-Injected and Forced Mouth Opening (FMO) Models. **(A)** Graph depicting the TMJ and orofacial withdrawal thresholds, as measured by von Frey filament, following CFA injection (n = 6 mice per group). (**B**) Graph depicting the TMJ and orofacial withdrawal thresholds determined by von Frey filament following FMO (n = 7 mice per group). BL: baseline, CFA: Complete Freund’s Adjuvant. Error bars indicate S.E.M. **p < 0.01; ***p < 0.001; two-tailed Student’s t test.

Peripheral sensitization of nociceptors usually takes two forms. One is a sustained spontaneous activity, and the other is a response to innocuous stimuli[8]. To assess whether CFA injection or FMO alters the spontaneous activity of TG neurons, we first observed the activity of TG neurons using *in vivo Pirt*-GCaMP3 Ca^2+^ imaging of intact TG. The total number of spontaneously activated neurons increased in both the CFA and FMO groups compared to the sham controls, and the increases were due to increases in two subgroups, small (<20 μm) and medium (20-25 μm) diameter neurons (Figs. 4A and B). We next asked whether TG neurons were sensitized in response to noxious or non-noxious stimuli. For this, we applied von Frey filament (mechanical), acetone (thermal), hot/cold water (thermal), or capsaicin (chemical) to the V3 region. Following application of 0.4 g filament to the V3 region, activation of small-diameter neurons but not of medium-diameter neurons was increased in CFA-injected mice, whereas activation of both small and medium-diameter neurons was increased in FMO mice (Figs. 5A and B). When we applied acetone to the V3 region of CFA-injected mice, the number of activated neurons was increased due to an increase in activation of small diameter neurons (Fig. 5C). Cold (4°C) water induced an increase in activated neurons of FMO mice (Fig. 5D), but hot (50°C) water did not induce any changes in activated neurons of V3 regions of either CFA-injected or FMO mice (Figs. 5E and F). Capsaicin injection into the V3 region increased the number of activated TG neurons in both CFA-injected and FMO mice (Figs. 5G and H). In CFA-injected mice, small and medium diameter neurons contributed to the increase of activated neurons, whereas in FMO mice, all three subgroups contributed to the capsaicin-induced increase (Figs. 5G and H). These data indicate that CFA injection and FMO differentially sensitize TG neurons to various stimuli and suggest that FMO causes behavioral and cellular changes related to TMD symptoms.

**Fig. 4.**
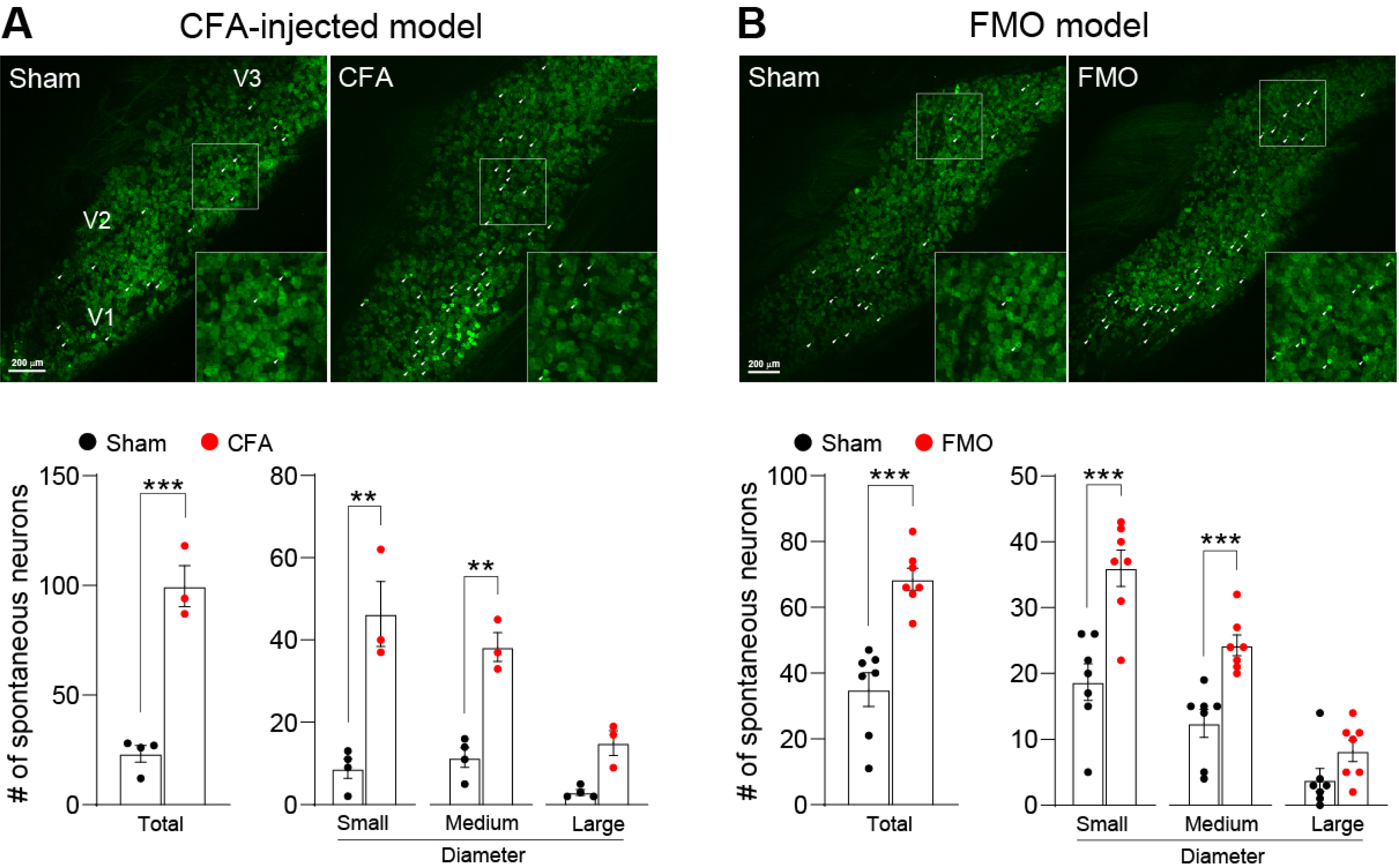
Spontaneous Activity increase of TG Neurons Following CFA-TMJ Injection or FMO. **(A)** (*Upper*) Representative images of spontaneous activities in *in vivo Pirt*-GCaMP3 Ca^2+^ imaging of intact TG in CFA-TMJ-injected mice. V1 (ophthalmic), V2 (maxillary), and V3 (mandibular) indicate the location of neuronal cell bodies in images of intact TG. White arrowheads indicate spontaneously activated neurons. (*Bottom*) Number of total, small, medium, and large spontaneously activated neurons from each group in CFA-TMJ-injected mice (n = 3 mice per group). (**B**) (*Upper*) Representative images of spontaneous activities in FMO mice. (*Bottom*) Number of total, small, medium, and large spontaneously activated neurons from each group in FMO mice (n = 7 mice per group). CFA: complete Freund’s adjuvant. FMO: forced mouth opening. Error bars indicate S.E.M. **p < 0.01; ***p < 0.001; two-tailed Student’s t test.

**Fig. 5.**
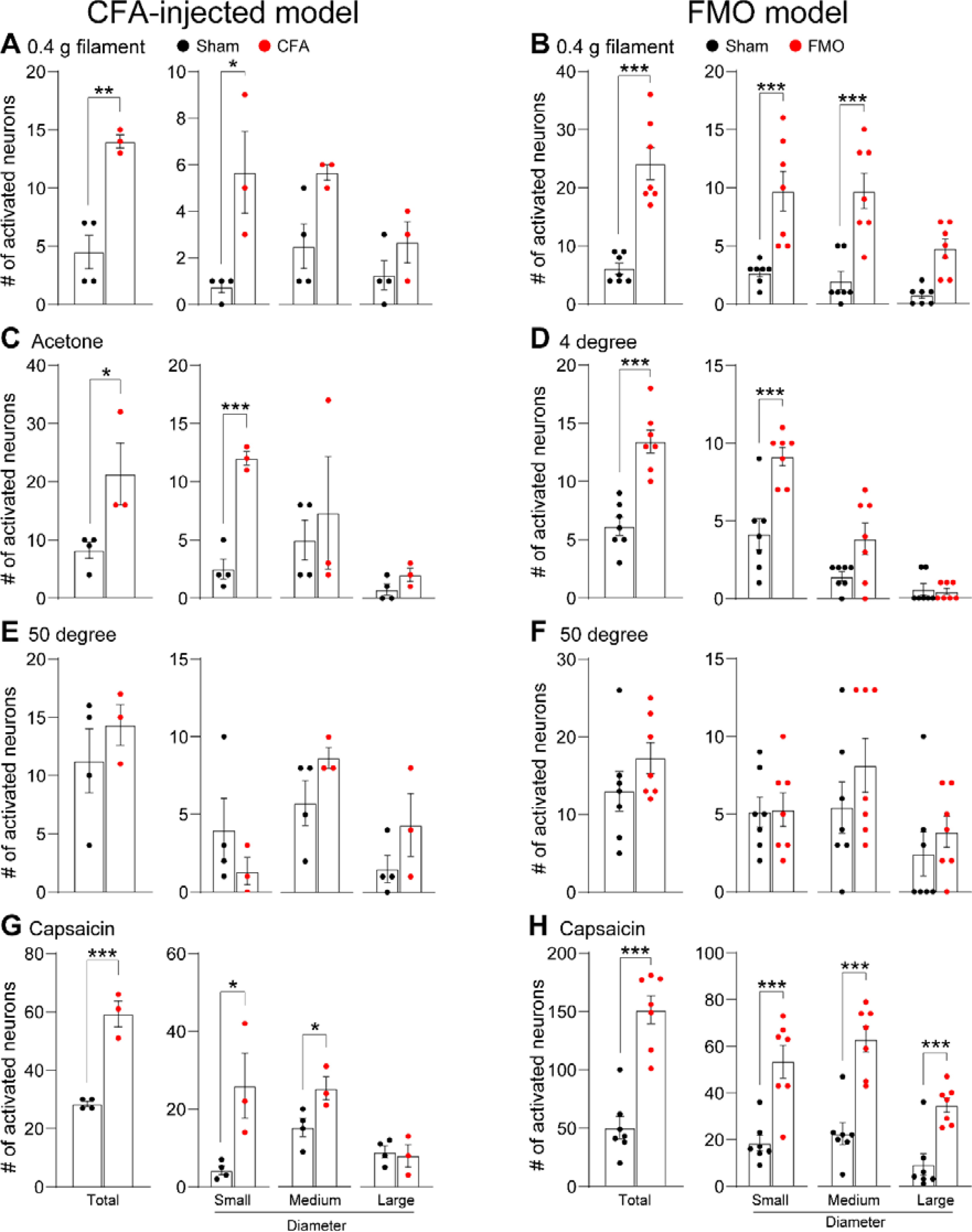
Differential Activation Patterns of TG Neurons in CFA-TMJ-Injected or FMO Mice in Response to Various Mechanical, Thermal, and Chemical Stimuli. Number of activated TG neurons in response to various stimuli in CFA-TMJ-injected (**A, C, E, and G**) (n = 3 mice per group) and FMO (**B, D, F, and H**) (n = 7 mice per group) mice. The graph represents the number of individual neurons activated by 0.4 g von Frey filament (A and B), Acetone (C), 4lJ water (D), 50lJ water (E and F), and capsaicin (G and H). CFA: Complete Freund’s Adjuvant. FMO: forced mouth opening. Error bars indicate S.E.M. *p < 0.05; **p < 0.01; ***p < 0.001; two-tailed Student’s t test.

### CGRP antagonist alleviates FMO-induced sensitization of TG neurons

CGRP levels are elevated in patients with TMD and are positively correlated with pain levels[1,2,16,21,29]. Thus, CGRP is thought to contribute to the pathogenesis of TMD, and indeed, CGRP antagonists have been shown to alleviate TMD symptoms in preclinical studies[5,17,25,32]. To determine if artificially increasing CGRP levels induced pain behavior, we administered CGRP directly into the TMJ and tested for pain behavior. CGRP injection induced hypersensitivity (Fig. 6A), which lasted up to 2 days after CGRP injection. Conversely, to determine whether a CGRP receptor antagonist (BIBN-4096) inhibited pain behavior in the FMO model, we induced mechanical hypersensitivity with FMO and then intraperitoneally injected a CGRP receptor antagonist. Hypersensitivity was gradually attenuated by successive injections (Fig. 6B). We used GCaMP3 Ca^2+^ imaging to determine whether the CGRP receptor antagonist attenuated FMO-induced sensitization of TG neurons. The antagonist was administered on day 4 after FMO under the same conditions as in Figure 6B, and GCaMP Ca^2+^ imaging was performed 2 hours later. We found that the increases in activated neurons seen previously (Figs. 4B and 5E-H) were attenuated by the CGRP antagonist (Figs. 6C-G). However, the increased number of activated neurons resulting from application of cold (4°C) water was not reduced by the CGRP antagonist (Fig. 6E). These results confirm the sensitization of TG neurons in the FMO-induced TMD model and indicate that GCaMP Ca^2+^ imaging enables mechanistic studies with various drugs and stimuli.

**Fig. 6.**
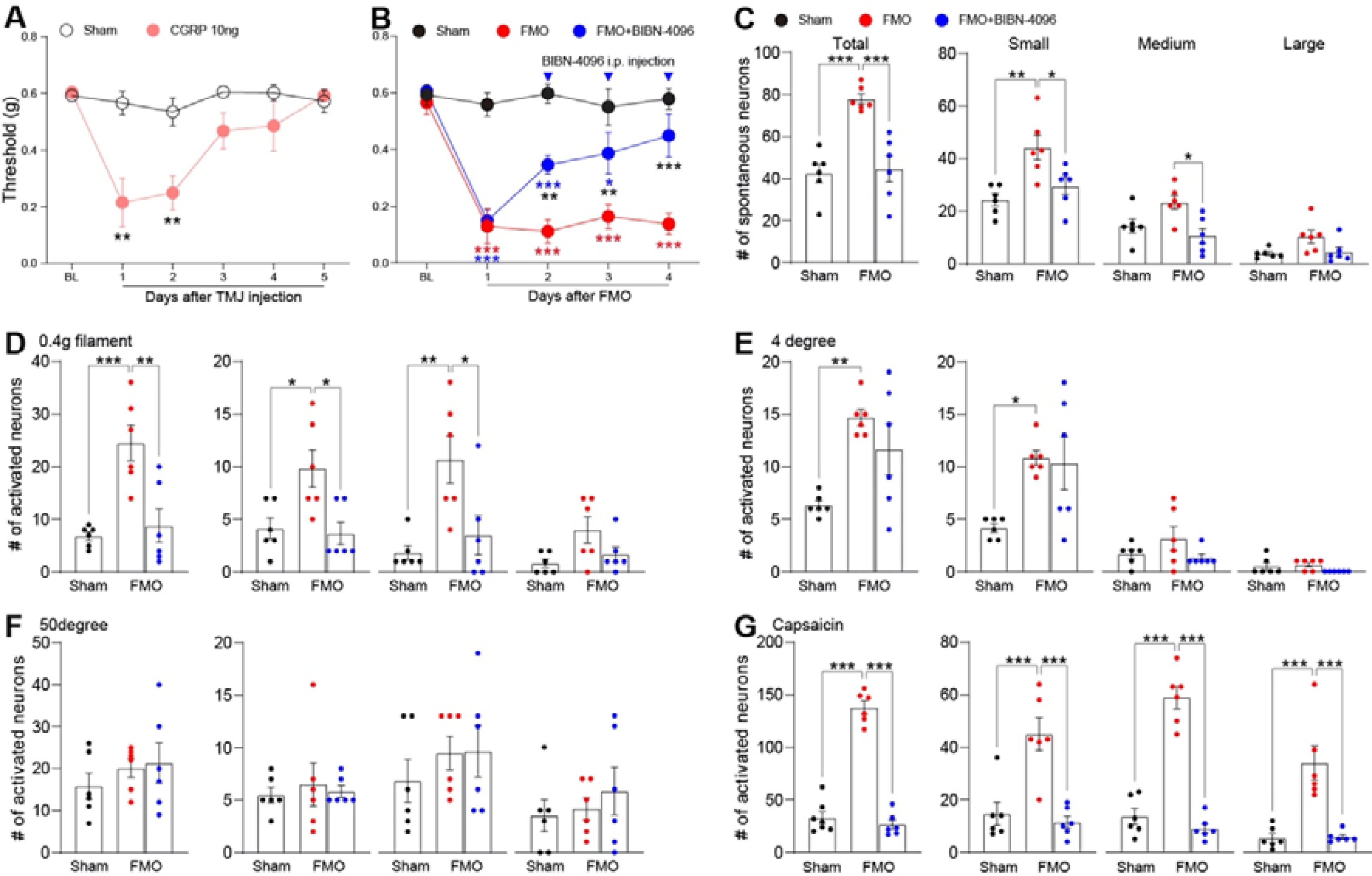
Evaluation of CGRP-Induced Hypersensitivity and Its Attenuation by a CGRP Receptor Antagonist in FMO Mice. **(A)** Graph showing head withdrawal thresholds tested by von Frey filament following CGRP-TMJ injection (n = 5 mice per group). (**B**) Head withdrawal thresholds observed in FMO mice treated with BIBN-4096 (CGRP receptor antagonist, 10 mg/kg i.p.) (n = 6 mice per group). (**C**) Number of total, small, medium, and large spontaneously activated neurons in FMO mice following BIBN-4096 treatment (n = 6 mice per group). (**D-G**) Number of individual TG neurons activated by 0.4 g von Frey filament (D), 4lJ water (E), 50lJ water (F), and capsaicin (G) following BIBN-4096 treatment in FMO mice. CGRP: calcitonin gene-related peptide. FMO: forced mouth opening. TMJ: temporomandibular joint. Veh: vehicle. Error bars indicate S.E.M. *p < 0.05; **p < 0.01; ***p < 0.001; (A) two-tailed Student’s t test; (B) two-way ANOVA with Tukey’s multiple comparison post-hoc test; (C, D, E, F, and G) one-way ANOVA with Tukey’s multiple comparison post-hoc test.

## Discussion

TMJ involves daily jaw movements, such as chewing, swallowing, speech, and other automatic movements, including yawning, grinding, or clenching. TMJ dysfunction in TMD patients can devastate daily life activities in humans[30]. However, available treatment methods and options only provide limited effectiveness. The unmet therapeutic needs of TMD patients stem from our poor understanding of the pathophysiological mechanisms of TMD. Here, we showed that *in vivo* GCaMP3 Ca^2+^ imaging of intact TG allows the simultaneous monitoring of nearly 3000 neurons in living mice and could be used to determine whether the sensitization of TG neurons occurs in clinically-relevant translational TMD animal models. Moreover, we confirmed that a CGRP receptor antagonist alleviates FMO-induced hypersensitivity of TG neurons. In addition, our use of *in vivo* GCaMP3 Ca^2+^ imaging of intact TG has provided nuanced insights into TG neuronal sensitization in TMD, laying a foundational step towards enhanced therapeutic strategies, as evidenced by the mitigating effects of CGRP antagonist on FMO-induced hypersensitivity.

The articular disc of TMJ has a limited ability to redistribute joint stress, making this joint susceptible to damage from overloading. Overloading causes disc displacement or joint degeneration, which leads to inflammation and the release of inflammatory mediators[6]. This inflammation can cause peripheral sensitization of nociceptors in the TMJ[23], ultimately resulting in pain and discomfort related to TMD[6]. Because of the role that inflammation plays in the development of TMD-related pain, CFA injection into TMJ in mice provides a reasonable TMD model. However, this model has limitations. CFA injections are limited in simulating the complexity of chronic inflammatory responses. While CFA can induce acute inflammatory reactions and histological modifications in the TMJ, including extracellular matrix accumulation and soft tissue alterations, CFA does not cause cartilage and bone degradation. Furthermore, this model is not fully representative of the underlying causes driving the sustained pain and degeneration experienced by human patients, as it does not encompass the unidentified foundational elements contributing to the persistence of these conditions[9]. Because excessive TMJ activity and prolonged mouth opening are risk factors in development of TMD[6], the FMO model of TMD-related pain is attractive[15,33]. FMO causes prolonged hypersensitivity in the V3 region of mice, which lasted for at least 32 days after FMO^9^, which is longer than the CFA-injection model. These characteristics suggest that FMO is sufficient to induce sensitization of TG neurons innervating TMJ.

Peripheral sensitization of the afferents originating from the TG significantly contributes to ongoing craniofacial pain[31]. Peripheral neuropeptides, receptors, and ion channels activating or sensitizing the nociceptive afferents have been identified[4,13]. TRPV1, TRPA1, and TRPV4 have been identified as contributing to the sensitization of TG in animal models of TMD[7,22,32]. However, direct studies on the sensitization of TG neurons in TMD animal models still need to be completed. Here, we monitored the spontaneous activity and response to stimulation of TG neurons in live mice in TMD animal models and showed that the neurons were more highly sensitized than in controls. We also confirmed that CGRP inhibitors attenuated FMO-induced TG sensitization. To our knowledge, this is the first time that the sensitization of TG neurons has been measured directly in a TMD model *in vivo*.

Neuropeptides are known to contribute to nociceptor sensitization[3,20]. In peripheral tissue, neuropeptides, including CGRP and substance P, are released from sensory neurons in a process of neurogenic inflammation and can activate their specific receptors to promote inflammation or sensitize nociceptors[24,26]. CGRP has been associated with craniofacial pain, including TMD[27,31]. In particular, in migraine, an increase in CGRP levels of peripheral tissues activates receptors on nociceptors and then sensitizes nociceptors[12]. New medications have been developed based on this mechanism and are now commercially available for migraine patients[11]. The effectiveness of these drugs in migraine continues to be confirmed[28]. In TMD, the relationship between CGRP levels and pain has also been confirmed[1,2,16,21,29], and preclinical studies have shown antinociceptive effects of CGRP antagonists[5,17,25,32]. However, the role of CGRP in TMD remains unclear. In this context, our confirmation of the effect of CGRP antagonists through *in vivo* GCaMP Ca^2+^ imaging represents a significant step forward in understanding CGRP-related mechanisms in TMD. As the systemic treatment of CGRP antagonists in this study did not completely exclude the involvement of the CNS, future studies are needed to investigate the mechanisms behind the effects of CGRP antagonists that we have identified.

This study confirmed that FMO, a novel TMD model, induces significant hypersensitivity in the V3 region, accompanied by TG sensitization, as confirmed by *in vivo* GCaMP3 Ca^2+^ imaging of intact TG. The finding reported here are the first confirmation of TG sensitization in an animal model of TMD and of the antinociceptive effect of a CGRP antagonist that attenuates TG sensitization, suggesting that *in vivo* GCaMP Ca^2+^ imaging of intact TG may provide the means to fully elucidate the pathogenesis of TMD and contribute to finding new therapeutic targets in preclinical studies.

## Data availability

Datasets analyzed during the current study are available from the corresponding author on reasonable request.

## Acknowledgements

This work was supported by National Institutes of Health grants NIDCR DE026677, DE031477, and NS128574 (Y.S.K.) and a Rising STAR Award (Y.S.K.) from the University of Texas System.

## Author contributions

Conceptualization: H.S., M.C. and Y.S.K. Resources and mouse strains: J.S. and R.G. Experimentation and acquisition of data: H.S., Z.Y., J.S., and R.G. Data analysis: H.S. Writing the manuscript: H.S., M.C. and Y.S.K. Funding acquisition and supervising project: Y.S.K.

